# Assessment of Auditable Pharmaceutical Transactions at Public Hospitals in Gamo Gofa, Southern Ethiopia, a Comparative cross-sectional study, July 2017

**DOI:** 10.1101/378000

**Authors:** Mende Mensa, Akililu Ayele, Biruk Wogayehu

## Abstract

**Background:** Availability of essential medicines is necessary to maintain health of the community. In Ethiopia, availability of medicines was low (65%), with high expiry rate (8.24%), low patient knowledge on correct dosage (50.5%) and satisfaction on pharmacy services (74.5%). To avert these problems, the government had endorsed legislation on a system called “Auditable Pharmaceutical Transactions and Services (APTS)”. However, the outcomes and challenges in implementation of this system were not assessed.

**Objective:** To assess the implementation status of APTS and its challenges at public hospitals in Gamo Gofa Zone Southern Ethiopia, April 2017.

**Methods:** Facility based Cross sectional study was conducted in two APTS implementing hospitals in Gamo Gofa zone. Semi structured Self-administered questionnaire was distributed to all pharmacy staffs in selected hospitals. APTS reports of 12 months (with different characteristics) were reviewed. Four hundred patients were interviewed by data collectors about patient knowledge and satisfaction using WHO questionnaire. The data were entered and analyzed using Statistical package for social science students/SPSS version 20. T-test and linear regression was used to evaluate significant differences between two hospitals with level of significance pre-set at p-value ≤0.05.

**Results:** All dispensing units in primary hospital had six (75%) out of the eight essential equipment for dispensing practice. it was found that respondents in general hospital stated higher scores in general setting of outpatient pharmacy (4.58 Versus 4.25; P <0.001), but lower scores for availability and cost of medicines (4.24 vs 4.43; P<0.05) when compared with those in primary level hospital. There was no significant difference in instruction of medicine provided by dispenser (2.58 vs 2.59; P>0.05), dispenser client interaction (3.09 vs 4.08; P>0.05) and total satisfaction score (2.09 vs 2.02; P>0.05).

**Conclusion and recommendations:** In our study Quality of Auditable Pharmaceutical Transactions and Service was low, especially regarding patient knowledge about medicines, unaffordability of medicines, less availability of prescribed drugs, poor transparency of pharmaceutical transactions, insufficient counseling practice and limited facilities for dispensing such as, key medicines, formularies and standard guidelines. We therefore recommend the following measures responsible bodies to improve these gaps y taking administrative actions and providing continued education and training for dispensers.

## Introduction

As per universal declaration of human rights by United Nations in 1948, “everyone has the right to a standard of living adequate for health and the right to security in the event of sickness or disability”. Medicines are vital instruments to complement healthcare service delivery for every citizen of the world. [1-3]. Ethiopia is currently on double burden of communicable diseases such as pneumonia, tuberculosis, HIV/AIDS and non-communicable diseases such as diabetes, hypertension and cancer [4]. To prevent and treat such diseases, huge capacities are needed including; health facilities, trained human power, availability of medicines and efficient utilization of resources [5, 6, 7]. In addition to this, factors that leads to wastage of medicines in health facilities such as; expiry, pilferage, theft and irrational use of medicines should be prevented. In turn, to perform the aforementioned duties, pharmacy organizations and patient flow within pharmacies should be properly addressed [8].

Globally more than 50% of all medicines are prescribed, dispensed or sold inappropriately, and half of all patients fail to take medicines correctly and hence the overuse, underuse or misuse of medicines harms people and waste resources. More than 50% of all countries do not implement basic policies to promote rational use of medicines. This is high in developing countries; only less than 40% of patients in the public sector and 30% in the private sector are treated according to clinical guidelines [9].

Pharmacy organization of health facilities, workflow within pharmacy outlets, the number, mix and ratio of pharmacist to client ratio are the basic elements to be fulfilled to deliver quality pharmacy services and attain appropriate patient satisfaction [8].

The pharmacies of hospitals in Ethiopia should be organized as outpatient, inpatient and emergency pharmacies and a central medical store of each directed by a registered pharmacist [10]. In addition, the hospital has to have adequate personnel, equipment, premises and facilities required to store pharmaceutical supplies and carry out compounding, dispensing and counseling activities. The work flow should be designed in such a way that customers should enter in one gate of the pharmacy outlets and exit in another, in a way inside the pharmacy; customers see prescription evaluator, biller, cashier, and medicines use counselor in a queue [8].

Transparency and accountability is another big challenge in the pharmaceutical sector. The World Bank has identified corruption as “the single greatest obstacle to social and economic development keeping millions of people trapped in poverty” and labeled a “cancer” [11].

The pharmaceutical sector is particularly vulnerable to corruption and unethical practices since the commercial reality of the pharmaceutical market tempts many different actors [8, 11]. As per the WHO strategy, improving good governance of pharmaceutical management in public health facilities is very important especially for disadvantaged, poor and vulnerable populations [11, 12]. The Federal Ministry of Health (FMOH) of Ethiopia had developed the Ethiopian Hospital Reform Implementation Guidelines (EHRIG) which includes the pharmacy service reforms [13].

Further to implementation of EHRIG in hospitals for the last five years, a system was designed that assumed to curtail the aforementioned pharmacy service drawbacks, called Auditable Pharmaceuticals Transactions and Services (APTS). The system, APTS is being put in to law regionally in Amhara 2011 [14], Dire dawa in 2012 [15], and SNNP in 2014 [16] and by the Federal Government [17, 18]. Many other regions have also drafted regulation to implement the system and FMOH of Ethiopia decided to be scaled up APTS nationwide.

APTS is a service delivery scheme that assumed to enables establishment of transparent and accountable medicines transaction and service provision. The ultimate objectives of APTS are to: institute ethical, transparent and responsible pharmacy practice that enables health facilities optimize utilization of medicines budget; improve access to medicines; continually improve the number, skill, mix & efficiency of pharmacy workforce, improve documentation and pharmacy premises and workflow, generate reliable and consistent information on products finance and services for decision making, improve patient knowledge on prescribed medicines and customers satisfaction. The system is intended to enables pharmaceutical transactions and service to be audited at any time [8].

APTS has five main pillars: Efficient budget utilization, transparent and accountable transactions, reliable information, effective workload analysis including; performance measurement and workforce deployment and improving customer satisfactions [8]. The APTS system [8] declares that there are many factors which affect the quality and volume of pharmacy service provision; including, lack of training that intern reflected by lack of knowledge and capacity, chaotic workflow, poor infrastructure, insufficient equipment and facilities needed to give the service, lack of using the highest efficient mix of services units of pharmacy, medicines budget and number of professionals.

Globally, in developing and industrialized countries alike, efforts to provide health care are facing new challenges. These include the rising costs of health care, limited financial resources, shortage of human resources, inefficient health systems, the huge burden of disease, and challenges to relate to treatment that one third of the world’s population does not yet have regular access to essential medicines [19, 20]. For many people, the affordability of medicines is a major constraint due to high price especially in private sector reaching in some cases 80 times the international reference price and requires over 15 days’ wages to purchase 30 days of treatment [19, 20, 21]. In low and middle income countries, because of high prices, medications account for 25% to 70% of total health care expenditures, compared to less than 15% in high-income countries. Inaccessibility and unaffordability to essential medicines are aggravated by medicines diversion from government to private, theft, non-transparency, nonsystematic selection, poor procurement and wastage due to expiry, irrational use, and poor pharmacy organization and workflow [8, 19, 20, 21].

A recent report of the President’s Malaria Initiative to Congress of the US Government indicated that until April 2014, the stealing is continuing and there was no solution solicited in Africa [22]. As per the study of World Bank in collaboration with anti-corruption authority of Ethiopia, even though corruption is uncommon compared to other African countries, pharmaceutical sector is found to be one of the two most corrupted sectors in Ethiopia that donated products are being diverted for private resale within Ethiopia and abroad [23, 24].

Studies showed that the root causes of drug diversion in Ethiopia includes: non-transparent transaction; while medicines entered in the store, issued to sections and dispensed to patients, patients used to buy medicines with a receipt prepared by a cashier who is unable to write the names and full descriptions of medicines. The type, quantity and price of medicines that are transacted had not been traced. Therefore, a system that can transparently show step by step flow of medicines until it reach the intended patient is becoming mandatory [8]. A recent baseline assessment for APTS implementation done by FMOH in collaboration USAID/SIAPS project, revealed that: patient knowledge on how to take their medicines; concerning dose, route of administration, frequency and duration showed that only 50.5% clients properly know all parameters [25].

In Ethiopia, various findings showed that essential medicines are poorly available (65%) [26], with high expiry rate (8.24% nationally) [27]. There are poor information on product and financial values of medicines, inefficient utilization of medicines budget, poor pharmacy infrastructure and chaotic work flow, all together resulting in poor quality of medicines management and erratic dispensing activities including counseling services and low overall patient satisfaction on pharmacy services (74.5%). [8, 13, 25]

In order to solve the aforementioned problems that the concept of APTS was innovated in Ethiopia, piloted in Amhara Region, Debre Markos Referral Hospital, in 2011 [8, 28], after comments are incorporated it is scaled up in health facilities throughout the country. There are few preliminary studies that assessed the outcomes of APTS have been documented by Amhara RHB and individual hospitals. However there is no similar study conducted in the study area. Therefore this study was conducted to evaluate APTS implementation status at general and district hospitals in Gamo Gofa zone, southern Ethiopia.

A study in Kenyatta National Hospital, Kenya, indicated that “low employee’s capacity, inadequate technology adoption for health service, ineffective communication channels and insufficient financial resources resulted to decrease in provision of health service quality [29]. A study conducted on factors influencing pharmacist performance in Great Britain, showed that; “ pharmacist performance is affected by characteristics such as age, gender, ethnicity, place of primary qualification, workplace factors, workload and mental and physical health problems, alcohol use or drug addictions” [30].

Workflow in pharmacy services is a problem in many African countries. Its inefficiency also has a negative impact in all over performance of the health facility. There are different models in work flow of pharmacy services like were “single server-multiple queue models and multiple servers with multiple queue models”. Finally, after staff re-orientation the streamline process, the best model that reduces waiting time from 167.0 to 55.1 minute which indicated a 67% reduction waiting time was adopted by consensus and practiced [42].

In the wall Street journal, a survey showed that antimalarial medicines are diverted from east to West Africa due to lack of transparency of medicines supply management system. Therefore, a system that can transparently show step by step flow of medicines until it reach the intended patient is becoming mandatory [31, 32].

Epidemiological study conducted in India showed that less than half reported that they did not ask and were not told how to store their medicines properly at home. Less than one third (30.4%) of study participants reported that they did not ask the doctor about any possible side effects of their medicines and more than two thirds (72.4%) discontinued their treatment course when they felt that their symptoms disappeared [33]. Another study conducted in Afghanistan showed that the patients who know all the seven WHO drug use indicators that enables on how to take dispensed medicines (the name, dose, route of administration, the frequency, duration, precaution, storage) ranged from less than 10% to 60% as shown in the graph below [34].

A study done in Kenya health facilities indicated that the incidence of expiry of medicines in dispensing shelves were found to 2.3% in government health facilities where as 1.9% in private health facilities [35]. Similarly study conducted in Uganda showed that high contribution of the expiry medicines to be due to storing medicines that treat rare diseases (81.8%) and drug donation (56 %) [36]. Similar study in Ethiopia revealed that the national averages expiry rate of medicines was found to be 8%, 2% and 3% in health facilities, regional drug stores and private drug retail outlets, respectively [25].

Study conducted on quality of hospital service in eastern Ethiopia showed that the percentage of patient satisfaction for pharmacy service was 65% being less than laboratory service (75%) [37]. Another study conducted in Ethiopia during collection of APTS baseline data indicated that there was an overall wastage of 3,281,562.20 ETB ($164,078.11) in 2012, accounting to an average of 3.9% of the total value of medicines received by 6 hospitals. In 2013, the value of wastage was estimated to be birr 10,684,221.09 indicating an average wastage rate of 8.3%; in 2014 total wastage of 1,542,491.6 ETB ($77,124.58) indicating an average of 5.1% wastage rate [25]. This rate of expiry was found to be equivalent to the rate of expiry of medicines taken during national HSDP-IV (2010-2015) baseline [27].

In the APTS baseline assessments conducted at different times in these hospitals, overall patient satisfactions on pharmacy services were found to be; 77% in Debre Markos Referral Hospital [38] and 40% in Felege Hiwot Referral Hospital [39]. In the APTS baseline assessment data showed that the patient knowledge on how to take dispensed medicines ranged that percentage of patients who knew all parameters were 15.5% whereas who knew all five basic drug use indicators(dose, route, name, frequency, storage and precaution) [38, 40, and 41].

### Conceptual framework for APTS evaluation; Adapted from Logic Model

**Figure 1:**
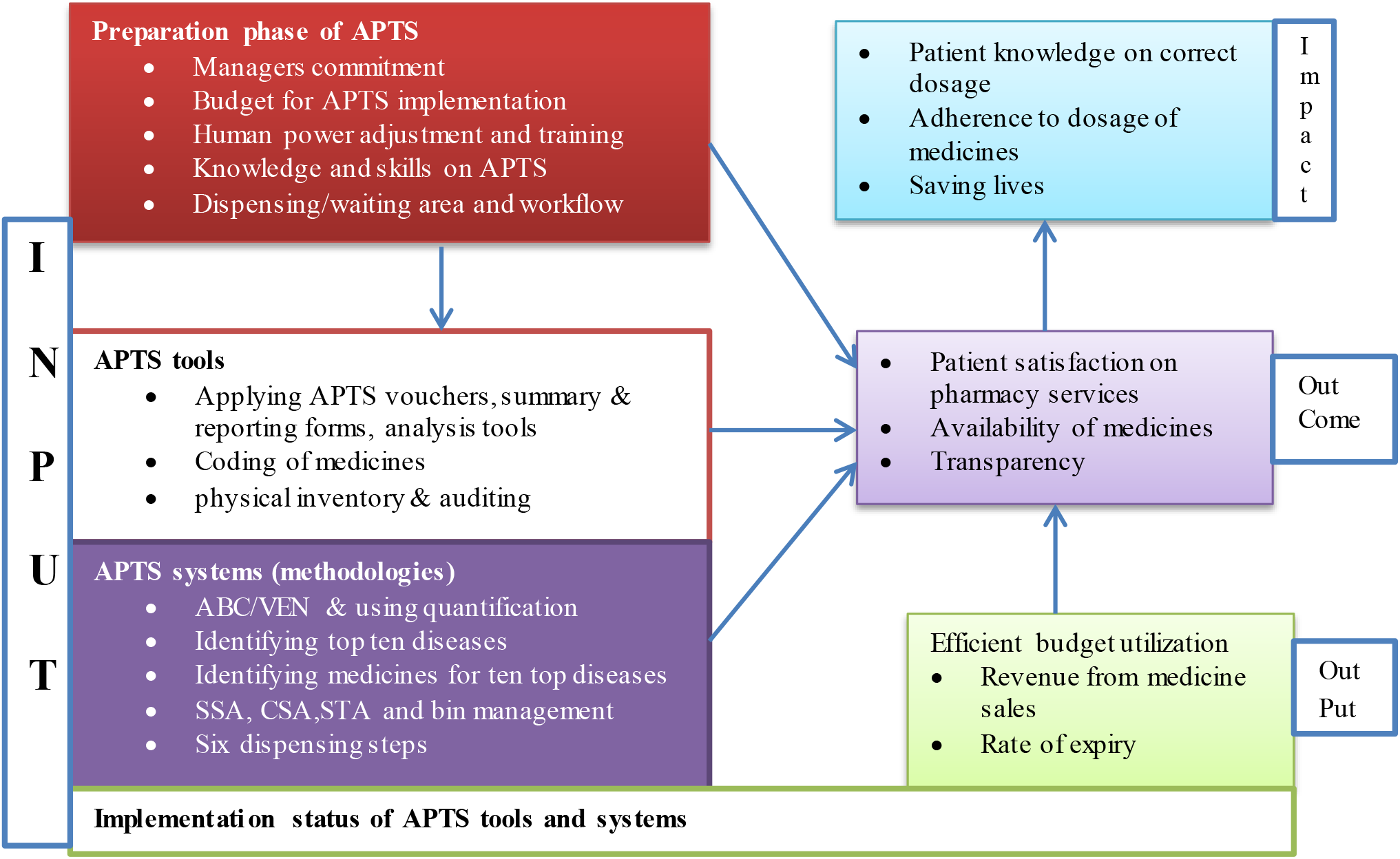
Conceptual framework for APTS evaluation; adapted from Logic Model Flowchart for Program Evaluation, March 2015

### Methodology

#### Study area and period

The study was conducted in hospitals from different woredas of Gamo Gamo Zone in July 2017. The Gamo Gofa zone has 15 woredas and one City Administrations with a recently estimated population of 1,597,767 (2007) people. The general elevation the Zone ranges from 600-3300 meter above sea level. The zone has a land mass of 12,581.4 square kilo meters. Gamo Gofa Zone has huge capacity of health service delivery system, focusing on prevention of diseases, with a capacity of Health Posts, 59 Health Centers and 4 Hospitals [Two general hospitals (Arbaminch and Sawula) and Two district hospitals (Chencha and Malo Laha)] [42]. Arba Minch General Hospital and Chencha District Hospital were randomly selected using random numbers from the list of general and district hospitals in the study area for this study.

### Study Design

Facility based Cross-sectional study design was conducted.

### Population

#### Source populations

The source populations to identify implementation status and challenges were all pharmacy staffs and CEOs of selected hospitals. The source populations for patient knowledge and satisfaction were all patients who got pharmacy service on the data collection period in the selected hospitals and the source populations for expiry rate, revenue from medicines sales, documents to be reviewed were monthly reports of APTS starting from the first APTS monthly report generated and submitted to RHB and or FMOH onwards, ABC/VEN analysis documents performed in the APTS implementation year/s and stock status analysis made in the same year/s.

#### Study populations

CEOs and head pharmacist of selected hospitals, all pharmacy and finance staffs working as cashiers, patients attending selected hospitals to receive pharmacy service during data collection, and monthly reports of APTS starting from the first months of APTS monthly report production on wards, sampled ABC/VEN analysis documents performed in the APTS implementation year/s and sampled stock status analysis made in the same year/s.

#### Inclusion criteria

- All pharmacy and finance staffs working in the pharmacy and having experience of at least six months
- Patients who received pharmacy service in the selected hospitals during the study period and willing to participate in the study with age greater than or equal to eighteen were taken.
- Monthly reports of APTS

#### Exclusion criteria

- Staffs who are in annual leave during the study period
- Staffs who are sick during the study period
- New staffs who were employed in less than six month period in the hospital
- Patients who were very sick and unable to give information and also who are not willing
- Documents which are disorganized

### Variables

#### Dependent variables

- APTS implementation status

#### Independent variables

- Patients socio demographic characteristics, Waiting time, Educational level of pharmacy professional’s, Pharmacy organization and workflow, Dispensing /waiting area that fulfill APTS standards, Dispensing counter and Seated service for special counseling in OPD and chronic care pharmacies

### Sample Size and Sampling Technique

#### Sample size determination

The sample size for APTS implementing status was all pharmacy staffs that fulfilled the inclusion criteria. For patient knowledge to dispensed medicines, availability of prescribed medicines and patient satisfaction on pharmacy services WHO recommends minimum of 100 samples for comparative analysis of drug use among facilities. The sample size for document review will be all APTS monthly reports produced from each hospital starting from the first month of APTS implementation onwards, all stock status analysis findings and ABC/VEN analysis conducted in the APTS implementation period. The sample sizes for the in-depth interview will be all CEOs and head pharmacists from selected hospitals.

#### Sampling Techniques

For APTS implementation status and challenges data were collected by self-administered questionnaire from all pharmacy and finance staffs in the selected hospitals. For expiry rate and revenue; data was collected from each selected hospitals by reviewing various data sources-APTS monthly reports starting from APTS implementation onwards. All ABC/VEN analysis documents and all stock status analysis documents analyzed during the APTS implementation periods will be also reviewed. For patient knowledge and satisfaction consecutive sampling was until attaining the desired sample size.

### Data collection tools and Techniques

#### Data collection tools

To identify the implementation status of various result areas of APTS questionnaire was developed by research team after evaluation of different literatures. The portion of validated and standardized WHO drug use indicator assessment tool, which had also been adapted by the Federal Ministry of Health and RHBs during baseline assessment of APTS, was adapted for the APTS context and was used to collect the data by exit interview from patients served in pharmacies of selected health facilities regarding patient satisfaction and knowledge on correct dosage. To collect secondary data from APTS monthly reports of each hospitals, and ABC/VEN and SSA documents, a checklist was prepared.

#### Data collection techniques

Twelve data collectors and two supervisors were recruited for data collection and training was given to data collectors by Principal investigators concerning the data collection tool, interviewing procedures, the sampling technique to follow, to review document and related ethical considerations. The principal investigators were overseeing the performance of each data collector’s daily, progress made and gave comments for each step. The collected data were sent through EMS to the principal researcher from each of data collectors.

#### Data Processing and Analysis

The process attributes for quality were measured by observation of the client-dispenser interaction during dispensing process. Technical and interpersonal aspects of process attributes were measured independently. Interpersonal aspects of process attributes were measured by observing prescription evaluation, medicine counseling, labeling of medicines and prescription registration. Dispensing time was used as a measure of the technical aspect of process attribute. The time spent on issues not connected to the medicine being dispensed was not considered. Chi-square tests were used to compare differences in proportion for interpersonal skill of process attributes. Independent sample t-tests were used to compare technical skill of process attributes. Percentages of drugs actually dispensed were used as a measure of the outcome attributes of quality. We used independent two-sample t test to compare the satisfaction scores of items, domains and the overall satisfaction score between two hospitals. Following, multiple linear regressions were employed to compare changes of satisfaction scores between the two hospitals by adjusting for respondents’ religion, age, health status, level of education, sex, payment status, and residence, number of visit, income, employment and marital status. Knowledge of prescribed medicine score was assessed seeing the response to nine questions namely the dosage of drug, name of the drug, form of administration, duration of treatment, any drug or food interaction, any adverse effect, storage condition, attitude when one or more dose missed and frequency of administration. Each item was weighted according to the significance to safe drug use. Crucial items for the client to take the medication received higher scores. If the client correctly knew the name of the drug, the route of administration, the dosage and its frequency, a total of 20 points were attributed. Ten points was attributed, if the client knew the duration of treatment, attitude when one or more dose missed, any interaction with foods or drugs and any adverse effect. When calculating the knowledge score for each item the total number of drugs that the individual client has been dispensed was also considered.

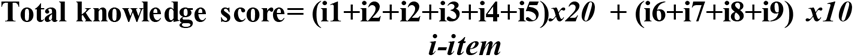

We employed independent two–sample t tests to compare score of items and total knowledge score between the two hospitals, while multiple linear regression analysis were then used to compare difference of total score between the two hospitals by adjusting for clients’ religion, age, health status, level of education, sex, payment status, residence, number of visit, income, employment and marital status.

Comparison of cost of treatment with the daily wage of the lowest-paid government worker (LPGW) is suggested by World Health Organization as a method of assessing medicine affordability. We assessed affordability of treating top ten diseases for adults and two common diseases for children by comparing total price of medicine at a standard dose to the daily wage of the LPGW of US $ *1.04*. We classified as a medicine unaffordable if it costs greater than a daily wage and affordable if it costs less than a daily wage.

Percentage of medicines actually dispensed is recommended by WHO as a means of assessing the degree of health facilities ability to provide the medicines which were prescribed. We assessed availability of prescribed medicine by asking clients number medicine prescribed and actual dispensed. Percentage of medicines actually dispensed was calculated by dividing number of medicines actually dispensed by the total number of medicines prescribed, multiplied by hundred. All analyses were conducted by using SPSS version 20.0

##### 4.8. Data Quality Management

To improve the consistency of the tools prepared, tools was originally prepared in English. To narrow language barriers during the interview, portion of tools that was used for patient exit interview was translated into local languages of the respective areas. The new portion of the tool was pretested in Nigisti Elleni Mohammed memorial Hospital to check whether the tool is sensitive enough to tempt interviewee in the intended manner and gather the necessary information needed. Data collector pharmacists were trained for one day by principal investigators. Final discussion was made with data collectors before and after the start of the assessment to make sure each member of the data collector fully understood the methods and tools. The guide was given for every data collector. Furthermore the principal researcher will oversee the whole data collection process. Once quantitative data is entered in to SPSS, all questioners were reviewed to ensure accuracy of data entry.

#### Operational Definition

- **APTS standard vouchers and sales tickets:** Models (19, 22), and sales tickets standardized by Federal Ministry of Finance for APTS implementation.
- **ABC analysis:** “A” class – 10 to 20 % of items that takes 70-80% of the overall total cost, “B” class – 10 to 20 % of items that takes 10-20% of the overall total cost and “C” class – 60-80% of items that takes 5-10% of the overall total cost
- **Affordability of medicine:** Affordability for a standard treatment of top ten diseases by comparing the total price of medicine at a standard dose according to Ethiopian standard treatment guideline to the daily wage of the lowest paid government unskilled employee at 20.5 Ethiopian birr (1.04 US $) per day. The cost of medicine for a full course of therapy for acute diseases and a 30-days’ supply of medicines for chronic diseases will be calculated and changed to the day wage. We will categorize as a medicine affordable “if it costs less than a day wage and unaffordable if it costs a day wage or more than a day wages”
- **APTS standard dispensing area and counter:** The dispensing areas of the outpatient and emergency pharmacies of a hospital that has entrance door, billing/prescription evaluation counter (with height 0.75cm for sitting service, 1.10 meter for standing service), for cashiers cubicle and medicines use counseling cubicle, and exit door in the opposite side of entrance.
- **Efficient budget utilization:** gain revenue from sales, rate of expiry less than 2% and affordability of medicines.
- **Implementation status of APTS tools and systems:** A hospital is said to be it has implemented certain result areas of APTS; if documents are found that showed the result area done as per the APTS guide for implementation or infrastructures are found being built. Example: availability of drug list, prioritizations of drug list by VEN, identifications drugs for ten top diseases, performing ABC analysis, conducting stock status analysis and taking interventions. Receiving, issuing, selling of medicines using vouchers/sales tickets approved by Federal ministry of finance, using drug codes, auditing reports, producing daily summary and monthly reports, dispensary has two doors, standard counters are built, man power adjusted, cashiers are inside the pharmacy, process are rearranged as per APTS guide etc.
- **Knowledge of professionals:** Level of understanding of the study participants (pharmacists, cashiers, accountants) about their assigned duties concerning APTS implementation is 100 % when they are asked to explain about their duties)
- **Key medicines:** Medicines used to treat 10 top disease are said to be key medicines
- **Management commitment:** defined as the devotion of managers of the hospital to allocate budget for renovation of dispensing area and employing human power.
- **Patient knowledge:** patients are considered that they know how to take their dispensed medicines if they answer at least all 5 basic WHO drug use indicators (the dose, route of administration, frequency, duration and storage) during exit interview.
- **Patient-days:** The number of days in which patients were served in a hospital
- **Patient Satisfaction:** patients are considered that they are satisfied if they answer either agree or strongly agree for the LIKERT scale questions and that will be re-coded in to new different variables
- **Revenue increment from sales of medicines:** revenue is increased if there is a positive slope of increment of revenue from sales of medicines starting from the baseline
- **Rate of expiry:** It is the percentage calculated by dividing the expired value in monetary forms to the stock available for sale.
- **Rate of sales of medicines:** Rate of sales of medicines is the percentage of sales of medicines divided by stock available for sale

#### Ethical Considerations

Ethical clearance was obtained from Arbaminch College of Health Sciences and Regional Health Bureau. A formal letter was written by Arbaminch College of Health Sciences, was submitted to the CEO of each hospital. Oral informed consent was obtained from each respondent for patient knowledge, availability of prescribed medicines and satisfaction prior to the interview. For the purpose of confidentiality and ethical issues, names of respondents from which information obtained were recorded and analyzed using uniquely identifying codes.

#### Dissemination of Results

The finding will be presented to the Arbaminch College of Health Sciences and other responsible bodies. Recommendations will be given to the Federal Ministry of Health, Regional Health Bureaus, respective Hospitals and relevant NGOs concerning the implementation of APTS. Finally attempt will be made to publish In peer reviewed national or international journal.

## Results

### Socio-demographic characteristics of respondents

We analyzed data of 400 respondents (200 from primary and 200 from general hospital). Participants differed, between the two hospitals, in age group, educational status, marital status and residence. More than 50% of the participants in both hospitals were males. More than one third of the participants in primary hospital had completed Primary school (33.5%) and more than one half in general hospitals had completed higher education (51.5%). With regard to marital status 164(82%) and 132(66%) were married in primary and general hospital respectively. Majority of the participants in both primary and general hospital were aged between 18 and 29 years (38% vs 39.5%). More than two third of the participants 191(95.5%) and 181(94.5%) in Primary and general hospitals pay for medicines respectively (Table 1).

### Structure attributes of quality: infrastructure, equipment, service, key medicines, guideline and format

All dispensing units in primary hospital had six (75%) out of the eight essential equipment for dispensing practice. These included the computer in pharmacy accountant office, scientific calculator, swivel chair, refrigerator, thermometer and tablet counter. Five (62.5%) out of eight of equipment required were available in the secondary hospital. Recent edition of APTS guideline, standard treatment guideline, good dispensing manual and drug formulary, procurement police manual were not available in both hospitals. With regard to pharmacy services, secondary hospital had eight (80%) out of the ten basic pharmacy services. These include Outpatient pharmacy service, Emergency pharmacy services, Inpatient pharmacy services, ART pharmacy, Clinical pharmacy services, Drug information services, Warehousing for medicines and Warehousing for medical supplies. However, only six (60%) out of ten basic pharmacy services were available in the primary hospital. All pharmacy accountant forms were available in the secondary hospital. However, only three (50%) out of six pharmacy accountant forms were available in the primary hospital (Figure 1).

**Table 1:**
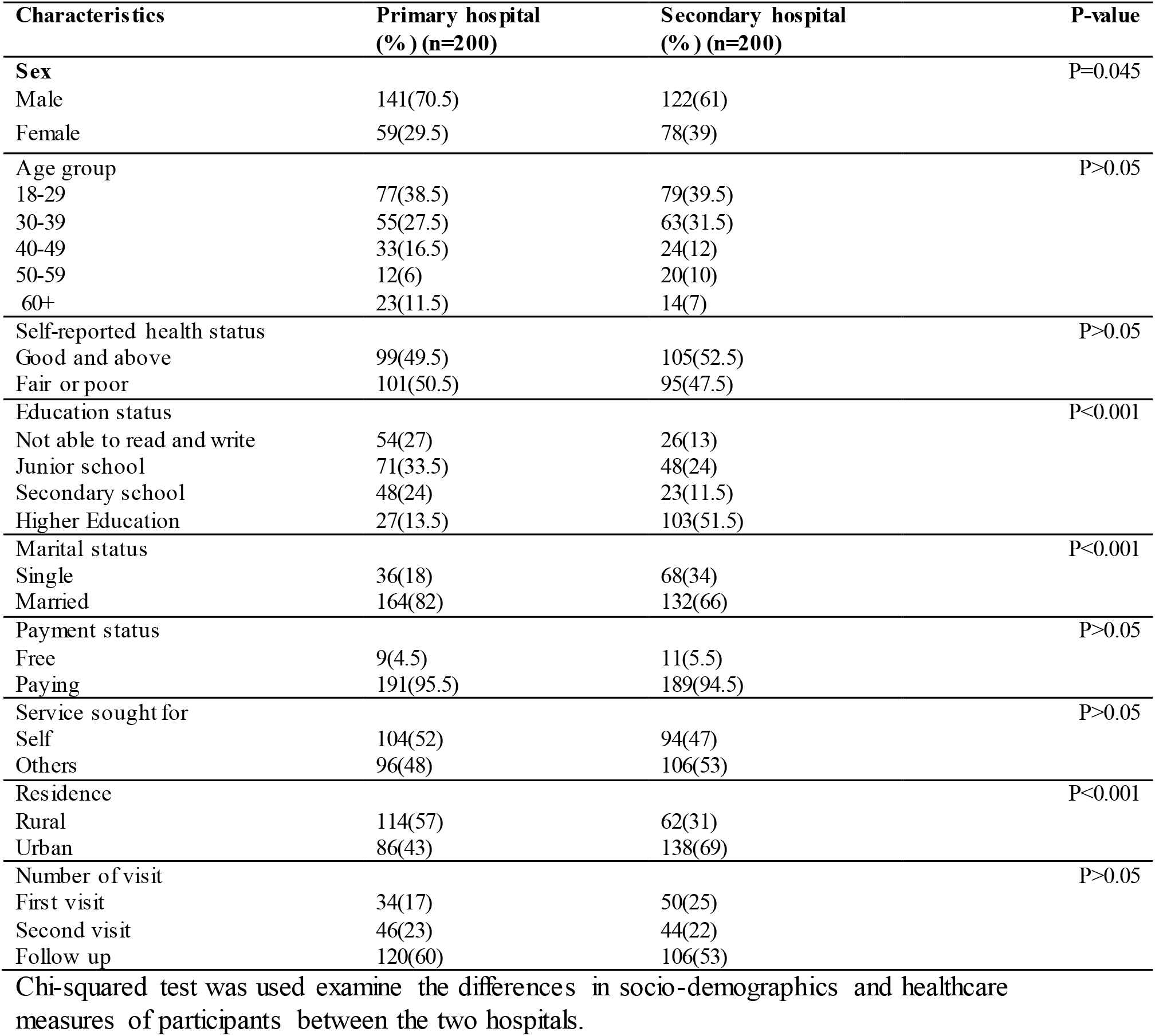
Socioeconomic and demographic characteristics and health care measures of the pharmacy service users in primary and secondary hospital.

| Characteristics | Primary hospital<br>(% ) (n=200) | Secondary hospital<br>(% ) (n=200) | P-value |
| --- | --- | --- | --- |
| **Sex** |  |  | P=0.045 |
| Male | 141(70.5) | 122(61) |  |
| Female | 59(29.5) | 78(39) |  |
| **Age group** |  |  | P>0.05 |
| 18-29 | 77(38.5) | 79(39.5) |  |
| 30-39 | 55(27.5) | 63(31.5) |  |
| 40-49 | 33(16.5) | 24(12) |  |
| 50-59 | 12(6) | 20(10) |  |
| 60+ | 23(11.5) | 14(7) |  |
| **Self-reported health status** |  |  | P>0.05 |
| Good and above | 99(49.5) | 105(52.5) |  |
| Fair or poor | 101(50.5) | 95(47.5) |  |
| **Education status** |  |  | P<0.001 |
| Not able to read and write | 54(27) | 26(13) |  |
| Junior school | 71(33.5) | 48(24) |  |
| Secondary school | 48(24) | 23(11.5) |  |
| Higher Education | 27(13.5) | 103(51.5) |  |
| **Marital status** |  |  | P<0.001 |
| Single | 36(18) | 68(34) |  |
| Married | 164(82) | 132(66) |  |
| **Payment status** |  |  | P>0.05 |
| Free | 9(4.5) | 11(5.5) |  |
| Paying | 191(95.5) | 189(94.5) |  |
| **Service sought for** |  |  | P>0.05 |
| Self | 104(52) | 94(47) |  |
| Others | 96(48) | 106(53) |  |
| **Residence** |  |  | P<0.001 |
| Rural | 114(57) | 62(31) |  |
| Urban | 86(43) | 138(69) |  |
| **Number of visit** |  |  | P>0.05 |
| First visit | 34(17) | 50(25) |  |
| Second visit | 46(23) | 44(22) |  |
| Follow up | 120(60) | 106(53) |  |

Chi-squared test was used examine the differences in socio-demographics and healthcare measures of participants between the two hospitals.

**Figure 2:**
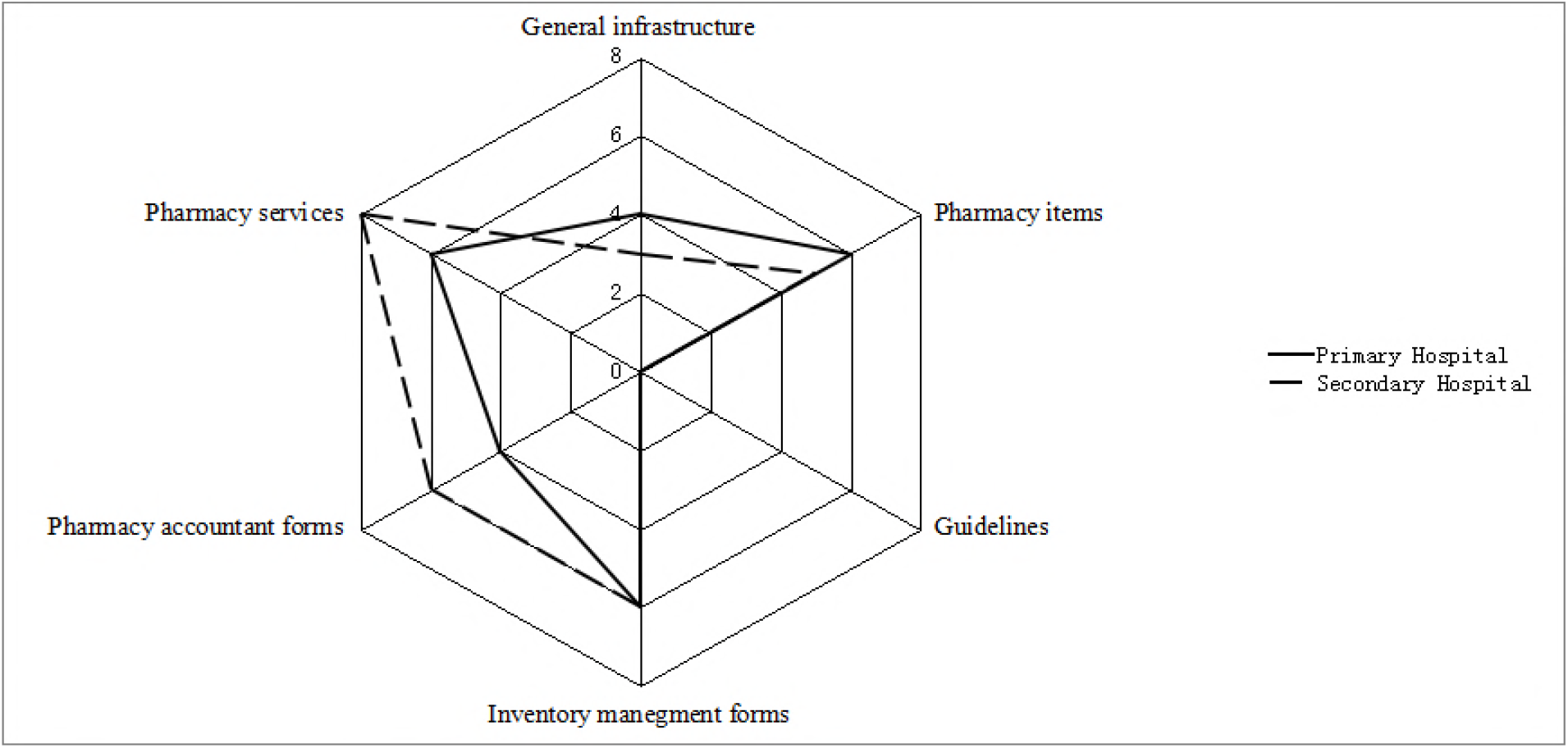
Infrastructure, equipment, service, guideline and format: structural attributes

### Process attributes of pharmacy service

There were significant differences in average scores observed for checking completeness of patient and prescriber information, with dispensers at general hospital scoring significantly higher than primary hospital dispensers(mean score 4.93 vs 4.37 and mean score 2.06 vs 1.13;p<0.001). In primary hospital more prescription evaluators were observed taking patient’s history (58.0% vs 21.5%; P<0.001) and writing the code of each billed drugs on sale tickets (100.0% vs 7%; P<0.001). General hospital appear to have higher scores for checking patient information and drug information on prescription (4.93 vs 4.37; P<0.001) than primary hospital. In general hospital, dispensers were more likely to explain the dosage form, purpose of the drug (22% vs 12.5%; P<0.05), when the drugs take (85.5% vs 13.5%) and importance of compliance (6.5% vs 2%; P<0.05) to the patient, but less likely to explain drug-food or drug-drug interaction, accurate dosing of the drug and storage condition of drug when compared with those in primary hospital. Dispensers in primary hospital were more likely to keep the privacy of the patients (98% vs 90.5%; P<0.01), make patient or relative to repeat the instructions and allow patients to ask questions (19.5% vs 5.5%; P<0.001) than those in general level hospital. More than 90% of the dispensed drugs in primary and secondary level hospital were not adequately labeled. The average dispensing time was significantly higher in primary hospital than general hospital (22.06 vs 14.93 seconds; P<0.01) (Table 2).

**Table 2:** Process attributes of pharmacy service at public hospitals in Gamo Gofa zone, southern Ethiopia, July 2017

| Characteristics c | Primary Hospital Dispensing practice(n=200) | Secondary Hospital Dispensing Practice(n=200) | Total (n=400) | $\rho$ values |
| --- | --- | --- | --- | --- |
| **Technical skills** |  |  |  |  |
| **Prescription Evaluation** | **SD** | **SD** | **SD** |  |
| Dispenser checks completeness of patient information (0-7) | 4.37(0 .48) | 4.93(0 .82) | 4.65(0.72) | 0.000 |
| Dispenser checks drug information (0-6) | 4.90( 0.38) | 5.01( 0.73) | 4.95(0.58) | 0.070 |
| Dispenser checks completeness of prescriber information (0-3) | 1.13( 0.33) | 2.06( 0.53) | 1.59(0.64) | 0.000 |
| **History taking** |  |  |  |  |
| Dispenser takes history of patient during evaluation of prescription (%) | 58.0 | 21.5 | 39.8 | 0.000 |
| Dispenser correctly code and bill price of drug | 100.0 | 7.0 | 53.5 | 0.000 |
| **Type of information given by the dispenser during counseling** |  |  |  |  |
| Dispenser tells patient name of the drug (%) | 0.0 | 1.5 | 0.8 | 0.248 |
| Dispenser tells patient dosage form of the drug (%) | 0.0 | 8.0 | 4.0 | 0.000 |
| Dispenser gives accurate information on the dosing of the drug (%) | 99.5 | 83.0 | 91.2 | 0.000 |
| Dispenser explains the purpose of drug (%) | 12.5 | 22.0 | 17.2 | 0.017 |
| Dispenser explains that when and how takes the drug (%) | 13.5 | 85.5 | 49.5 | 0.000 |
| Dispenser tells patient the duration of medication (%) | 13.0 | 8.5 | 10.8 | 0.197 |
| Dispenser gives information on drug-food or drug -drug interaction (%) | 43.5 | 21.0 | 32.2 | 0.000 |
| Dispenser gives information on possible side effects (%) | 0.0 | 0.5 | 0.2 | 0.990 |
| Dispenser gives information on contraindications (%) | 0.5 | 0.5 | 0.5 | 0.998 |
| Dispenser tells patients how to store drug(%) | 6.0 | 1.0 | 3.5 | 0.014 |
| Dispenser explains that importance of compliance (%) | 2.0 | 6.5 | 4.2 | 0.047 |
| Satisfactory counseling (%) | 0.0% | 1.5% | 0.8% | 0.248* |
| **Type of information labeled** |  |  |  |  |
| Dispenser labels patient name (%) | 0.5 | 2.0 | 1.2 | 0.372 |
| Dispenser labels drug name (%) | 4.5 | 4.0 | 4.2 |  |
| Dispenser labels when the drug should be taken (%) | 7.5 | 3.0 | 5.2 | 0.073 |
| **Prescribed drug or its equivalent is given(%)** | 99.5 | 95.0 | 97.2 | 0.014 |
| **Interpersonal skill** |  |  |  |  |
| Privacy ensured during counseling (%) | 98.5 | 90.0 | 94.2 | 0.001 |
| Integrity, fluency and configuration of speech (%) | 98.5 | 89.5 | 94.0 | 0.000 |
| Dispenser asks patients if she/he had any questions or concerns (%) | 28.0 | 10.0 | 19.0 | 0.000 |
| Dispenser makes patient or relative to repeat the instructions (%) | 19.5 | 5.5 | 12.5 | 0.000 |
| **Counseling duration(in seconds)** | 22.06(24.96) | 14.93( 18.23) | 18.49(22.12) | 0.001 |

\*Fisher exact test

### Availability of key medicines

The percentage availability of key medicines in primary level hospital was 60% compared to 80% in secondary level hospital (Table 3).

**Table 3:** Availability of key medicines at public hospitals in Gamo Gofa zone, southern Ethiopia, July 2017

| No. | List of key medicines | Primary Hospital | General Hospital |
| --- | --- | --- | --- |
| 1 | Amoxicillin with or without clavulanic acid | ✓ | ✓ |
| 2 | Oral Rehydration Salts | ✓ | ✓ |
| 3 | Artemether + lumefantrine (coartem) tabs | ✓ | ✓ |
| 4 | Mebendazole Tablets | ✓ | ✓ |
| 5 | Tetracycline Eye Ointment | – | ✓ |
| 6 | Paracetamol tablet/suspension | ✓ | ✓ |
| 7 | Rifampicin/Isoniazid/Pyrazinamide/Ethambutol | ✓ | ✓ |
| 8 | Medroxyprogesterone (depo) injection | ✓ | ✓ |
| 9 | Ergometrine Maleate Injection/Tablets | – | – |
| 10 | Ferrous Sulphate plus Folic Acid | – | – |
| 11 | Pentavalent DPT-Hep-Hib Vaccine | – | ✓ |
| 12 | Lidocaine injection | ✓ | ✓ |
| 13 | TAT injection | – | – |
| 14 | Diclophenac injection | – | ✓ |
| 15 | Doxycycline capsule | ✓ | ✓ |
| 16 | Cimetidine injection | – | ✓ |
| 17 | Ceftriaxone injection | ✓ | ✓ |
| 18 | Fluconazole tablet/capsule | – | ✓ |
| 19 | Ciprofloxacin tablet | ✓ | ✓ |
| 20 | Cotrimoxazole tablet | ✓ | ✓ |
| 21 | Metronidazole injection | – | – |
| 22 | Adrenaline injection | ✓ | ✓ |
| 23 | Ringer lactate solution | ✓ | ✓ |
| 24 | Glucose 40% solution | ✓ | ✓ |
| 25 | AZT /3TC/NVP | ✓ | ✓ |
| 26 | Benzyl penicillin Na | – | ✓ |
| 27 | Gentamycine | ✓ | – |
| 28 | Cloxacillin | ✓ | ✓ |
| 29 | Albendazole | – | – |
| 30 | Chloramphenicol eye drop | – | ✓ |
| Availability of key medicines (%) |  | 60% | 80% |

### Outcome of care

Respondents in a general hospital stated higher scores in general setting of outpatient pharmacy (4.58 vs 4.25; P <0.001), but lower scores for availability and cost of medicines (4.24 vs 4.43; P<0.05) when compared with those in primary hospital. The largest differences between the two hospitals were detected in general pharmacy setting (0.378) and availability and cost of medicines (0.179). Patients in a general hospital were more likely to receive labeled drugs than those in a primary level hospital (2.61vs 2.18; P<0.05). In terms of dispenser client interaction, more respondents in a primary hospital than in general hospital reported that their dispensers understood what they asked and their questions were answered in ways they understood (3.77 vs 3.20; P<0.05) (Table 4).

**Table 4:** Mean score of the selected items under each domain reported by respondents in primary and secondary level care

| Items | PCOP (SD) | SCOP (SD) | Unadjusted *p*-value | Adjusted health facility level effect $\beta$ (95% CI) |
| --- | --- | --- | --- | --- |
| **General setting of OPD** |  |  |  |  |
| Information for location of the pharmacy | 4.35(0.96) | 4.66(0.86) | 0.001 | 0.208(0.003,0.412) |
| Reference to other location | 4.42(0.82) | 4.75(0.72) | 0.000 | 0.328(0.154,0.503) |
| Space | 3.92(1.30) | 4.80(0.69) | 0.000 | 0.939(0.705,1.172) |
| Cleanness | 4.39(1.05) | 4.06(1.39) | 0.007 | -0.300(-0.575,-0.024) |
| Privacy | 4.24(0.93) | 4.35(1.00) | 0.256 | 0.213(-0.001,0.427) |
| **Availability and cost of medicines** |  |  |  |  |
| Cost | 4.44(0.86) | 4.16(1.10) | 0.006 | -0.314(-0.537,-0.090) |
| Availability | 4.39(1.05) | 4.05(1.38) | 0.007 | -0.300(-0.575,-0.024) |
| **Medicine information** |  |  |  |  |
| Labeling | 2.18(1.37) | 2.61(1.59) | 0.004 | 0.352(0.015,0.689) |
| Information for storage | 2.36(1.29) | 2.23(1.36) | 0.329 | -0.088(-0.390,0.213) |
| Side effect | 1.93(1.10) | 1.76(1.05) | 0.126 | -0.139(-0.382,0.105) |
| How to take | 4.40(0.76) | 4.41(1.08) | 0.956 | 0.014(-0.196,0.223) |
| Contraindication | 2.08(1.2) | 1.91(1.18) | 0.166 | -0.104(-0.374,0.165) |
| **Dispenser client interaction** |  |  |  |  |
| Feedback | 3.77 (1.40) | 3.20(1.77) | 0.000 | -0.471(-0.840,-0.103) |
| Waiting time during billing | 4.25(0.99) | 4.26(1.14) | 0.962 | 0.033(-0.190,0.256) |
| Time during counseling | 4.07(1.15) | 3.91(1.36) | 0.205 | 0.051(-0.235,0.337) |
| Professionalism | 3.90(1.12) | 3.92(1.17) | 0.896 | 0.038(-0.221,0.296) |
| Respect | 4.39(0.90) | 4.43(1.03) | 0.718 | 0.098(-0.117,0.313) |

### Patient satisfaction

The patients were found to be more satisfied in primary hospital than in general hospital about the cost of drugs (4.44 vs 4.16, P<0.01) and cleanness of dispensing room (4.39 vs 4.06; P<0.05) (Table 5).

**Table 5:** Comparison of mean satisfaction scores according to the level of health facility (n=400)

| Items | PCOP<br>(SD) | SCOP<br>(SD) | Unadjusted<br>*p*-value | Adjusted health facility level<br>effect $\beta$ (95% CI) |
| --- | --- | --- | --- | --- |
| General setting of OPD pharmacy | 4.25(0.69) | 4.58 (0.58) | **0.000** | **0.378 (0.234,0.521)** |
| Medicine information provided by dispenser | 2.59(0.78) | 2.58(0.77) | 0.949 | -0.055(-0.234, 0.125) |
| Dispenser client interaction | 4.08(0.71) | 3.94 (0.87) | 0.084 | 0.007(-0.169, 0.183) |
| Availability and cost of medicines | 4.43(0.59) | 4.24(0.71) | **0.005** | **-0.179(-0.327,-0.031)** |
| Total satisfaction score | 15.34(2.02) | 15.35(2.09) | 0.964 | 0.151(-0.316, 0.617) |

More than 70% and above of the respondents in primary and general hospital reported the scores of general setting of OPD pharmacy (81% vs 94.5%), dispenser client interaction (70% vs 69%) and availability and cost of drugs (84.5% vs 72%) above the point of reference of 3.75 (Figure 3).

**Figure 3:**
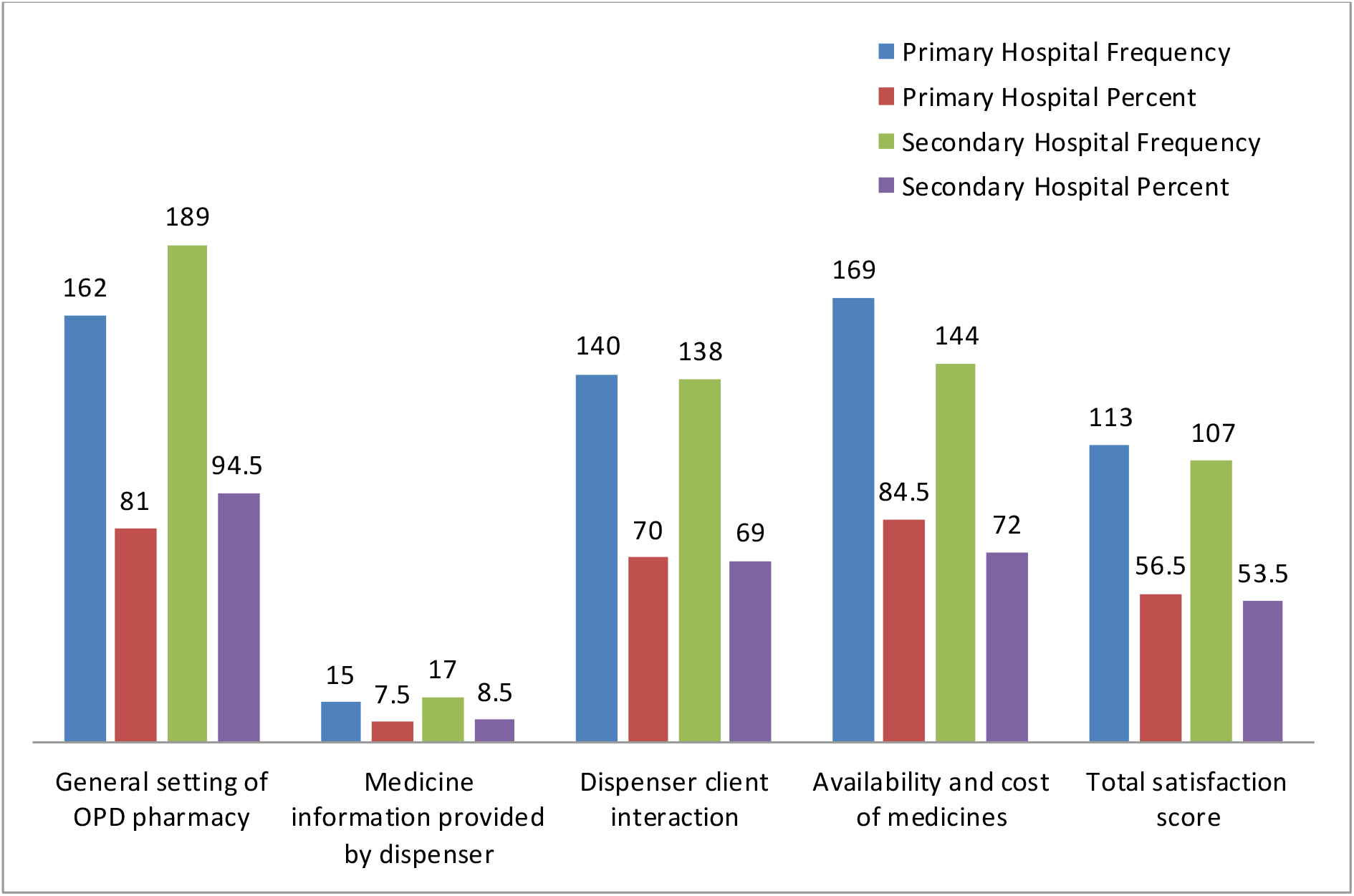
Percentage of the rating scores under good quality standard level by hospital type (Good quality ≥3.75 for each attributes and ≥15 for total score).

### Availability of prescribed drugs

Availability of all prescribed drugs was reported more frequently in primary hospital than in general hospital (90% vs 80%; P<0.01).Overall, 85% of the patients reported that all prescribed drugs were available in both hospitals. The percentage of drugs actual dispensed was 86.8% in the secondary hospital and 94.6% in primary hospital (figure 4).

**Figure 4:**
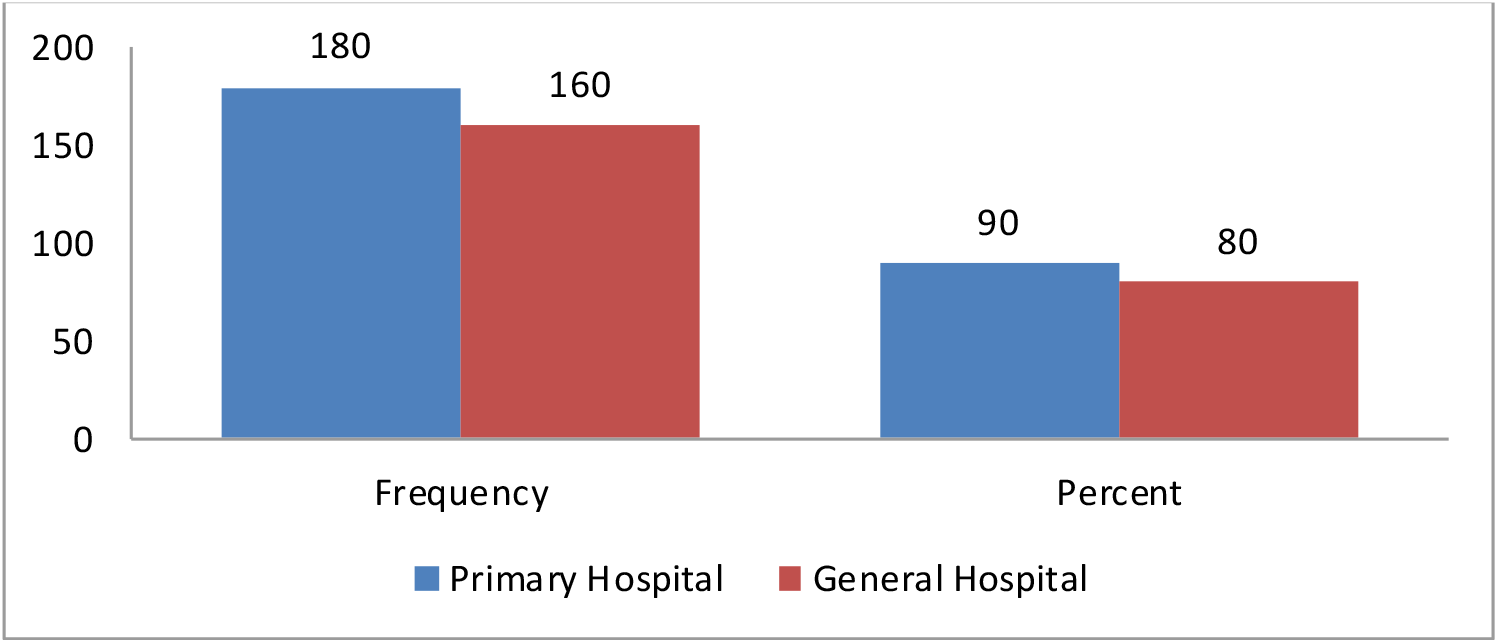
Availability of prescribed drugs at public hospitals in Gamo Gofa Zone, southern Ethiopia, July 2017.

### Knowledge of prescribed medicines

Patients were interviewed about the drugs at their concerning; name of drugs, dose, route of administration, frequency, duration of treatment, storage conditions and adverse effects, What to do when doses are missed and drug interactions with drug or foods. Less than one third 18 (26.86%) and 22(22.68%) of patients who received on drug do not know the name of the drug in Primary and General Hospital respectively. Less than one quarter patients received two drugs 7(9.45%), 12(16.21%) in primary hospital knew one drug and two drugs respectively. Concerning patients received two drugs from General hospital 16(21.33%), 18(24.0%) knew one drug and two drugs respectively. Regarding the dose of drugs 49(71.13%), 93(95.87%) Patients received one drug knew the dose of a drug. One (1.35%), 54 (72.97%) received two drugs from primary hospital, 11 (14.67%), 60 (80.0%) received two drugs from general hospital knew dose of one drug and two drugs respectively (Table 6).

**Table 6:** Frequency distribution of patient knowledge about dispensed drugs at Public hospitals in Gamo Gofa Zone, Southern Ethiopia, July 2017

| Knowledge about... | Number of drugs dispensed | Do not know | | Knows one drug | | Know two drugs | | Know three drugs | | Knows ≥ four drug | |
| --- | --- | --- | --- | --- | --- | --- | --- | --- | --- | --- | --- |
|  |  | PH | GH | PH | GH | PH | GH | PH | GH | PH | GH |
| Name of drug | 1 | 49 | 75 | 18 | 22 | 0 | 0 | 0 | 0 |  | 0 |
|  | 2 | 55 | 41 | 7 | 16 | 12 | 18 | 0 | 0 |  | 0 |
|  | 3 | 37 | 12 | 2 | 6 | 2 | 1 | 6 | 2 |  | 0 |
| | $\geq 4$ | 11 | 2 | 1 | 1 | 0 | 3 | 0 | 0 | | 1 |
| Dose of drug | 1 | 18 | 4 | 49 | 93 | 0 | 0 | 0 | 0 | 0 | 0 |
|  | 2 | 19 | 4 | 1 | 11 | 54 | 60 | 0 | 0 | 0 | 0 |
|  | 3 | 14 | 1 | 0 | 1 | 5 | 2 | 18 | 17 | 0 | 0 |
| | $\geq 4$ | 4 | 0 | 0 | 0 | 1 | 0 | 1 | 0 | 6 | 7 |
| Route of drug administration | 1 | 1 | 1 | 65 | 96 | 1 | 0 | 0 | 0 | 0 | 0 |
|  | 2 | 1 | 1 | 1 | 1 | 72 | 73 | 0 | 0 | 0 | 0 |
|  | 3 | 0 | 1 | 1 | 1 | 2 | 2 | 44 | 17 | 0 | 0 |
| | $\geq 4$ | 0 | 0 | 0 | 0 | 1 | 0 | 1 | 0 | 10 | 7 |
| Frequency of administration | 1 | 8 | 4 | 58 | 93 | 1 | 0 | 0 | 0 | 0 | 0 |
|  | 2 | 5 | 4 | 4 | 7 | 65 | 64 | 0 | 0 | 0 | 0 |
|  | 3 | 5 | 1 | 1 | 1 | 7 | 2 | 34 | 17 | 0 | 0 |
| | $\geq 4$ | 0 | 0 | 0 | 0 | 1 | 0 | 2 | 1 | 8 | 7 |
| Duration of treatment | 1 | 16 | 27 | 50 | 70 | 1 | 0 | 0 | 0 | 0 | 0 |
|  | 2 | 16 | 17 | 7 | 11 | 51 | 47 | 0 | 0 | 0 | 0 |
|  | 3 | 15 | 6 | 2 | 0 | 6 | 3 | 24 | 12 | 0 | 0 |
| | $\geq 4$ | 3 | 1 | 1 | 0 | 1 | 2 | 1 | 1 | 6 | 3 |
| Storage of drug | 1 | 22 | 49 | 44 | 48 | 1 | 0 | 0 | 0 | 0 | 0 |
|  | 2 | 28 | 38 | 1 | 1 | 45 | 36 | 0 | 0 | 0 | 0 |
|  | 3 | 12 | 10 | 0 | 1 | 2 | 0 | 33 | 10 | 0 | 0 |
| | $\geq 4$ | 4 | 2 | 0 | 0 | 0 | 0 | 1 | 1 | 7 | 4 |
| Adverse effect of drug | 1 | 62 | 76 | 5 | 21 | 0 | 0 | 0 | 0 |  |  |
|  | 2 | 69 | 60 | 1 | 3 | 4 | 12 | 0 | 0 |  |  |
|  | 3 | 43 | 17 | 1 | 2 | 1 | 0 | 2 | 2 |  |  |
| | $\geq 4$ | 12 | 4 | 0 | 3 | 0 | 0 | 0 | 0 | | |
| What do when one or more doses missed | 1 | 43 | 66 | 24 | 30 | 0 | 0 | 0 | 0 | 0 | 0 |
|  | 2 | 47 | 50 | 1 | 1 | 26 | 24 | 0 | 0 | 0 | 0 |
|  | 3 | 30 | 15 | 1 | 3 | 2 | 1 | 14 | 2 | 0 | 0 |
| | $\geq 4$ | 9 | 5 | 0 | 1 | 1 | 0 | 2 | 0 | | 1 |
| Drug interaction with drug or food | 1 | 50 | 51 | 17 | 45 | 0 | 0 | 0 | 0 | 0 | 0 |
|  | 2 | 51 | 31 | 6 | 8 | 17 | 36 | 0 | 0 | 0 | 0 |
|  | 3 | 29 | 10 | 3 | 1 | 4 | 4 | 11 | 6 | 0 | 0 |
| | $\geq 4$ | 7 | 3 | 0 | 1 | 1 | 0 | 2 | 1 | 1 | 2 |

**Note:** **DH:** Primary Hospital; **GH:** General Hospital

Finally Independent two-sample t test was employed to compare score of items and total knowledge score between the two hospitals. Linear regression analysis was used to compare difference of total score between the two hospitals after adjusting for confounders, there was no significant difference in the total drug knowledge score (89.38 vs 83.82; P>0.05) between the two hospitals (Table 7).

**Table 7:** Total knowledge score about dispensed drugs at Public hospitals in Gamo Gofa Zone, Southern Ethiopia, July 2017

| Hospital type | Mean(SD) | Unadjusted p-value | Adjusted health facility effect $\beta$(95% CI) |
| --- | --- | --- | --- |
| Primary level | 83.82(25.94) | 0.032 | 0.871(-4.793,6.536) ,P- 0.763 |
| Secondary level | 89.38(25.58) |  |  |

### Affordability of medicines

With regard to affordability of medicines most of the lowest priced generics needed to treat common uncomplicated conditions cost less than a days’ wages in the primary and secondary level hospital. The unaffordability of lowest priced medicines in the secondary hospital varies from 2.36 to 7.21 days’ wages. The unaffordability of the lowest priced medicines varies from 2.49 to 3.15 days’ wages in the primary hospital. The most unaffordable standard treatment was treatment of peptic ulcer with Amoxicillin 1 g tablet +Clarithromycin 500mg tablet+ Omeprazole 20mg tablet in both hospitals (days’ wages=3.15 vs 7.21) (Table 8).

**Table 8:**
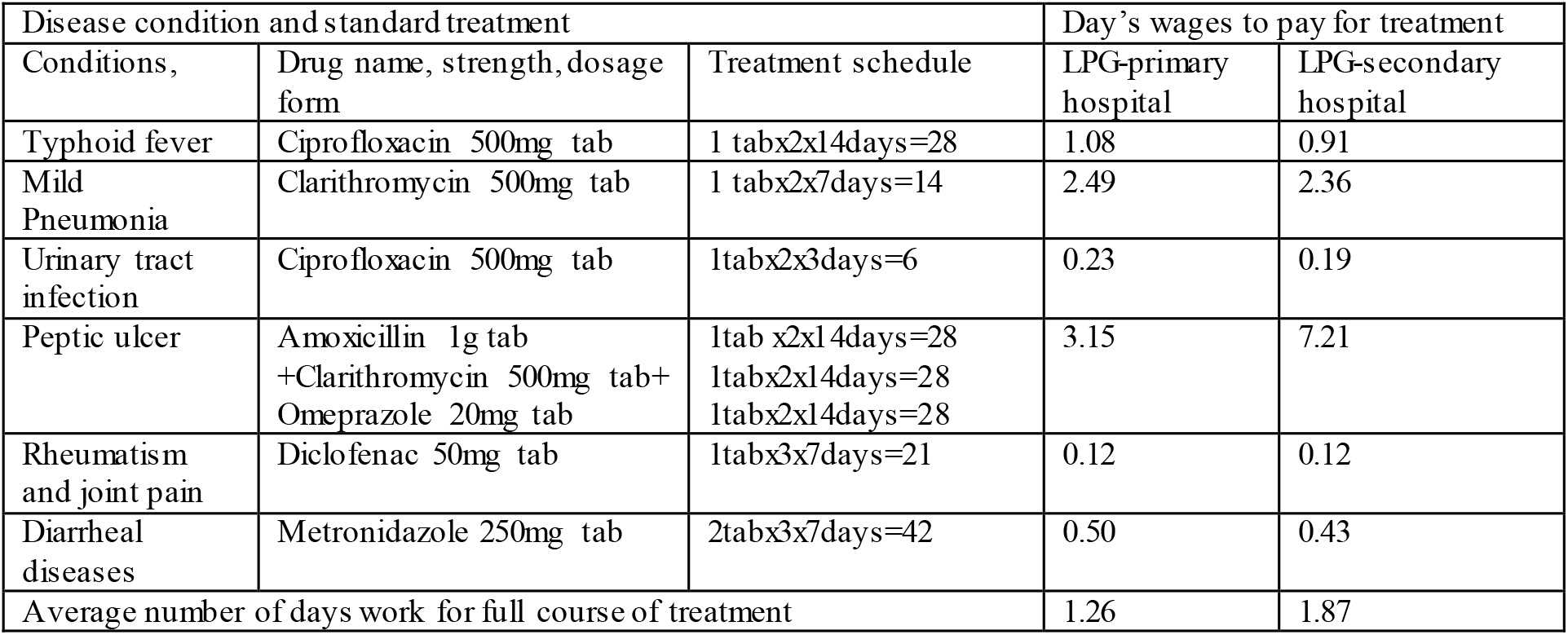
Affordability of medicines for common uncomplicated diseases at Public hospitals in Gamo Gofa Zone, Southern Ethiopia, July 2017

| Disease condition and standard treatment |  |  | Day's wages to pay for treatment |  |
| --- | --- | --- | --- | --- |
| Conditions, | Drug name, strength, dosage form | Treatment schedule | LPG-primary hospital | LPG-secondary hospital |
| Typhoid fever | Ciprofloxacin 500mg tab | 1 tabx2x14days=28 | 1.08 | 0.91 |
| Mild Pneumonia | Clarithromycin 500mg tab | 1 tabx2x7days=14 | 2.49 | 2.36 |
| Urinary tract infection | Ciprofloxacin 500mg tab | 1tabx2x3days=6 | 0.23 | 0.19 |
| Peptic ulcer | Amoxicillin 1g tab<br>+Clarithromycin 500mg tab+<br>Omeprazole 20mg tab | 1tab x2x14days=28<br>1tabx2x14days=28<br>1tabx2x14days=28 | 3.15 | 7.21 |
| Rheumatism and joint pain | Diclofenac 50mg tab | 1tabx3x7days=21 | 0.12 | 0.12 |
| Diarrheal diseases | Metronidazole 250mg tab | 2tabx3x7days=42 | 0.50 | 0.43 |
| Average number of days work for full course of treatment |  |  | 1.26 | 1.87 |

### Transparency and accountability of pharmaceutical transactions

With regard to transparency and accountability of the pharmaceutical transactions, the secondary hospital had only nine (45%) out twenty elements for making pharmaceutical transactions transparent and accountable. This included all medicines received in the store uses standard APTS receiving vouchers, all medicines issued uses standard APTS issuing vouchers, all medicines received in the last and recent transaction of the receiving model labeled with costs or estimated monetary values, all medicines issued in the last and recent transaction of the issuing model labeled with prices or estimated monetary values, the expiry dates of all medicines received by the recent transaction recorded in the receiving voucher, the batch numbers of all medicines received by the recent transaction recorded in the receiving voucher, and the hospital uses standard APTS cash sales tickets for cash transactions. Similarly the primary hospital had only ten out (50%) twenty elements for making the transaction transparent and accountable. In both hospitals, daily and monthly finance and service reports were not prepared in the last month. Service auditing, finance auditing and physical inventory were performed in both hospitals.

### Revenue from medicines

Beginning from the baseline revenue from medicines, putting into practice of APTS, the slope of the cash sales of medicines in primary hospital was found to be positive. However, revenue from medicines in the secondary hospital was found to negative slope. In primary hospital, revenue from medicines highly increased during implementation of APTS than in the secondary hospital (Figure 5).

**Figure 5:**
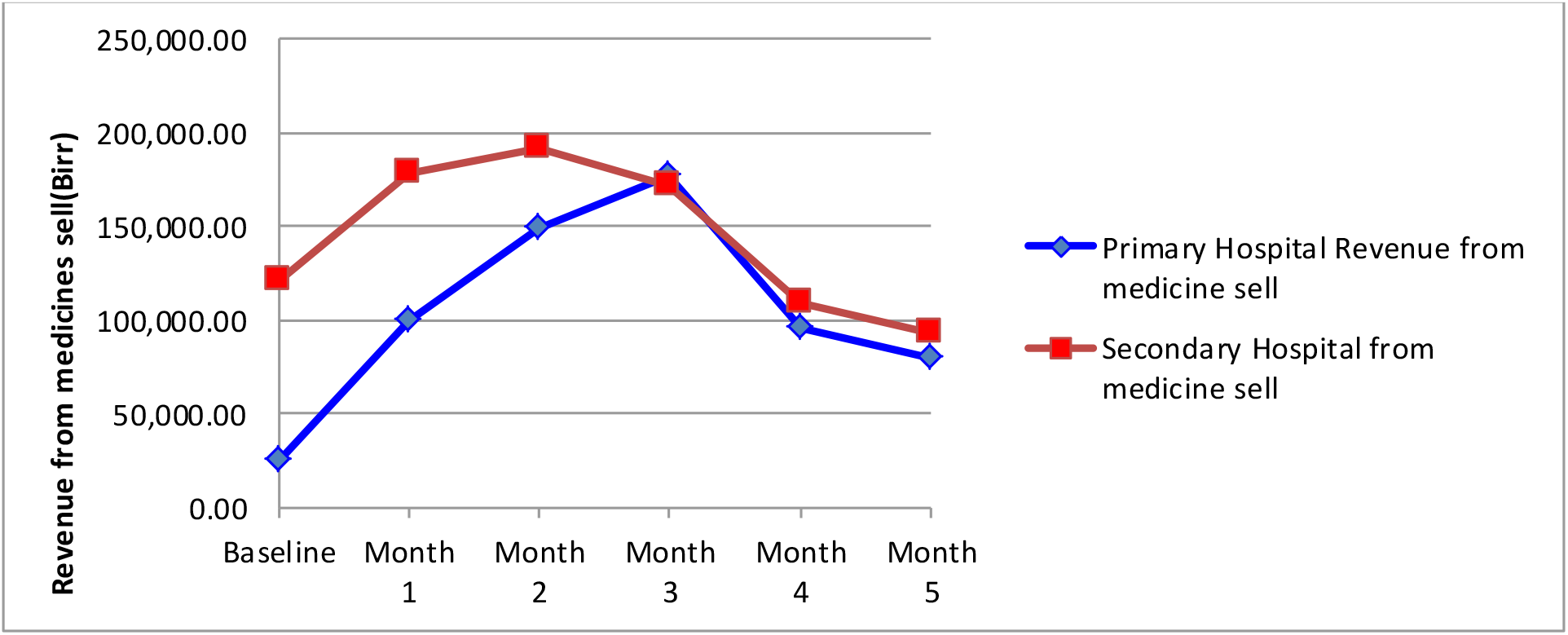
Trend of revenue from medicines from baseline (before APTS implementation) to successive five months at Public hospitals in Gamo Gofa Zone, Southern Ethiopia, July 2017

### Barriers to APTS implementation

In the primary hospital, the main barriers to APTS implantation included “high patient load (100%)”, “lack of training (83.3%)”, “lack of supervision” (66.7%) and “patient factors” (66.7%). In the secondary hospital, the main barriers included “high patient load” (100%),” lack of training” (66.7%) and lack of indemnity(33.3%).Overall, the main challenges for implementation of APTS involved “high patient load”(100%), “lack of training”(72.2%) and “shortage of human power”(66.7%). Lack of indemnity, lack of devotion of interest, lack of time, shortage of cashier, shortage of prescription papers and lack of good communication with prescriber and patient factors (patients have low attitude towards pharmacy, patients do not need to stay more time in the pharmacy) were also other barriers assessed(Figure 6).

**Figure 6:**
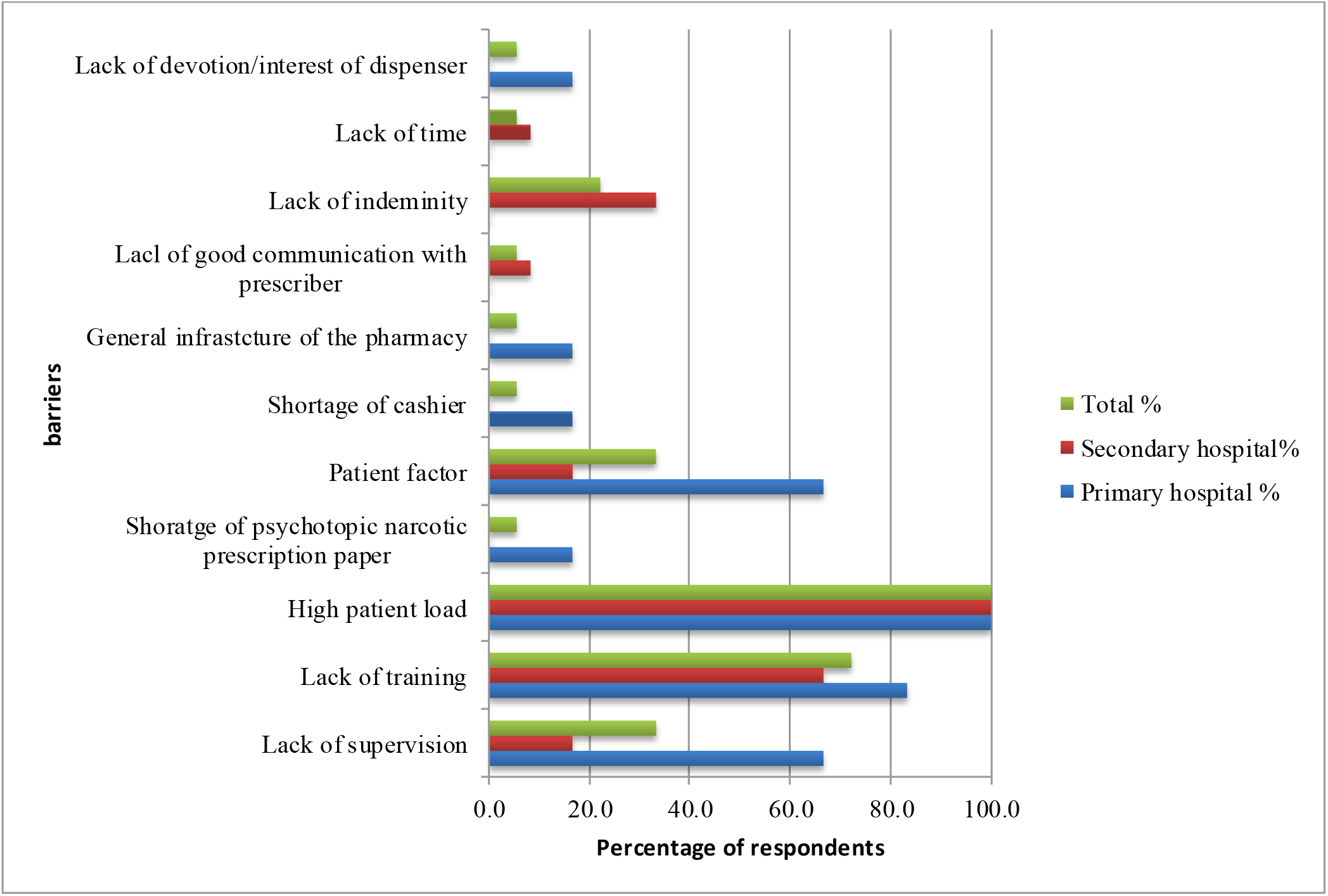
Dispensers responses to perceived barriers to APTS implementation at Public hospitals in Gamo Gofa Zone, Southern Ethiopia, July 2017.

## Discussion

This study revealed implementation status of APTS tools and systems and challenges at general district hospital. In this study the average dispensing time (22.06 and 14.93 seconds) in primary and general hospital respectively. A statistically significant difference exists between the results obtained in primary and in generally hospital on average dispensing time (p<0.01). The possible reason for this variation can be due to difference in man power, patient overload and set up of dispensary area. The average dispensing time in both hospitals was considered to be short(less than 3 minutes as per the WHO standard) [43]. Our finding is comparable to the finding from Jimma university specialized hospital (22.5 seconds) [44] and Kebrebeyah refugee health center (27.6seconds) [45]. Similar insufficiency of dispensing time was also reported in Tanzania (77.8seconds) [46], Nigeria (12.5seconds) [47], Mozambique (37seconds) [48], Swaziland (18.1seconds) [49] and Bangladesh (23seconds) [50], Jordan (28.8seconds) [51] and brazil (17seconds) [52].

Since pharmacists are the last healthcare personnel who see the patients or relatives before they take their drugs, dispensers need to provide adequate drug information to ensure the patients will all right and properly take their medications [53-55]. The duration found in both hospitals does not allow for inclusion of important drug related information such as drug regiment, importance of compliance, potential side effects, interaction with other medications and appropriate drug storage. Inadequate information about therapy could lead to non-adherence and consequent adverse drug events [56].

The present study revealed that, 0.0% and 2% of dispensed drugs were appropriately labeled in primary and general hospital respectively. This percentage values are lower than the recommended WHO value (100%). WHO recommends that each medicine label should include the patient name, drug regimen and drug dose [43]. The potential reason for this low value can be due to shortage of package for labeling, marker, attitude of dispensers and high patient load. Studies have shown that even in countries with advanced labeling practices, only 50% of drugs are taken as intended. Therefore this problem is more severe in countries with poor labeling practice. Our value is lower than reported in Eastern Ethiopia (64%) [57], Southwest Ethiopia (70%) [58], Bole Hora hospital, Southern Ethiopia (12.3%) [59], Tanzania (20.1%) [60] and Swaziland (55.9%) [49] but in line with the study conducted in selected public hospitals of Eastern Ethiopia(3.3%)[61],Nepal(1.4%)[62], Pakistan(6%)[63] and a study conducted in Cambodia (0.0%) [64].

History taking from patients that guide dispensing and counseling decisions such as identify concomitant medicines (being used currently used or stopped with in the last 21 days),alcohol use, adherence problem, renal or liver disease, pregnancy or breast feeding, allergy or any adverse drug reaction and ability to pay was done better in the primary hospital than general hospital.

The result of this study also showed that correctly billing and coding of prescribed medicines were done more better in the primary hospital than secondary hospital (100% vs 7%, p,<0.001). It implies that the primary hospital gives pharmacy accountants a better chance to carry out activities like preparing daily summary, monthly financial and service reports that are essential for making pharmaceutical transaction transparent and accountable. The information given in primary and in secondary hospital was the name of the drug (0.0%, 1.5%), dosage of the drug(0.0%, 8.0%), purpose of the drug(12.5%, 22%), dosing of the drug (99.5%, 83.5%), duration of medication (13%, 8.5%), possible side effects (0.0, 0.5%), contraindications (0.5%, 0.5%), storage condition (6%, 1%) and importance of compliance (2%, 6.5%) respectively. These findings are consistent with the results of another study where 20%, 3.8%, 100%, 8.9%, 0.0%, 0.0%, 0.0%, 0.0%, 0.0% and 7.7% of the patients were received counseling on the dosage form of the drug, purpose of the drug, dosing of the drug, duration of medication, drug interaction, possible side effects, contraindication, storage condition and importance of compliance of the, respectively [65].

In this study revealed that 0.0% and 1.5% of patients received satisfactory counseling at dispensing encounters in primary and secondary hospital, respectively. This finding is lower than the finding from Bahir dar city (32.8%) [66], South west Ethiopia (38.8%) [58], Botswana (91%) [67] and Pakistan (3.1%) [63]. World Health Organization’s drug use indictors stated that the percentage of satisfactory counseling on dispensed medicines should be 100%. However, in the present study the status of satisfactory counseling is still very low compared to the standard value. This deviation from the optimal value might be ascribed to shortage of counselors’, over - load of patients and counselors’ fail to practice good dispensing principles, medication counseling guide and code of ethics. Pharmacists or dispensers’ are the last health care providers with who a patient comes in contact before taking a drug. Additionally, dispensers are accessible to patients, often seeing them on several occasions between routine prescriber visits.

Our study revealed that when compared with those in primary hospital, patients in general hospital experienced better general setting of dispensing area, but lesser availability of prescribed medicines. Respondents in both hospitals had poor experiences in medicine information provided by dispenser but good experience in general setting of the pharmacy, interaction with dispenser and availability of prescribed medicines.

Patients in general hospital reported a higher score in general setting of the pharmacy than those in primary hospital, which indicated a better dispensing area (sufficient space, easily accessible reference to other location and clear information for the location of the pharmacy) in secondary hospital than that in primary hospital. Both respondents of the two hospitals, no significant difference were identified in total satisfaction score. This study found that patient satisfaction with labeling of medicines, information for storage, information on possible side effects and contraindications in both primary and secondary were too low, which is in line with a study conducted in a university hospital in northwestern Ethiopia [68] which found that the majority of patients were dissatisfied with labeling of medicine, information on possible side effects, storage condition and contraindication.

These findings are also consistent with the results of another study conducted in Saudi Arabia [69] and King Fahd Armed Forces Hospital [70]. In the pharmacy waiting time, cleanness, location reference to other services and counseling time domain our results better than results of a study conducted in Gonder university hospital [68]. High costs of medicines are major challenges to accessing medicines and getting better health outcomes. Affordability in this study is assessed in terms of the number of days the lowest paid unskilled governmental worker would have to work to pay for treatment course. At the time of the survey, the lowest paid unskilled governmental worker earned---Ethiopian birr (US$) as at 2017. On the average, the lowest paid government worker needed a 1.27 and 1.87 day’s wages to treat common disease conditions in primary hospital and secondary hospital, respectively. Least priced drugs are unaffordable for half of standard treatments of prevalent diseases in both hospitals since they cost more days’ wages for lowest paid unskilled governmental worker. Our finding is consistent with study of affordability in western part of Ethiopia [71], Ghana [72], Haiti [73], Brazil [74] and Guatemala [75].

Although the lowest paid employee wage was used as a measure of affordability it is likely that a significant part of the population earns less. In Ethiopia, significant numbers of population live below the poverty line. For this people, any out-of-pocket expense to drugs could be catastrophic.

In the present study, 94.6% and 86.8% of all medicines prescribed were provided in the primary and general hospital respectively. This variation might be partly ascribed to difference in the inventory management. This finding is lower than the finding from eastern Ethiopia (75.77%), (60.3%) [45] and the average of 12 countries (89%). This finding indicated that patients were prone to unnecessary medication charge by private pharmacies where profit of margin might reach more than 100% [76]. This study identified significant difference in total medicine knowledge score between the two hospitals.

### Strength and limitation

#### Strength of the study

- Adequate sample size use, Pre-tested tools were used and Data quality was maintained

#### Limitation of the study

- Being a cross sectional study, it doesn’t determine cause and effect.
- The responses might be influenced by socially desirable bias.
- The patient knowledge about dispensed drugs and satisfaction by pharmacy service was studied among outpatient pharmacy at the primary and secondary hospital and this finding of this study may not generalized to other departments

## Conclusion

This study revealed suboptimal quality of Auditable Pharmaceutical Transactions and Service quality in two hospitals, especially as it regards unsatisfactory knowledge score of medicines, unaffordability of medicines, less availability of prescribed drugs, poor transparency of pharmaceutical transactions, insufficient counseling practice and limited facilities for dispensing such as, key medicines, formularies and standard guidelines.

### Recommendations

Based on the findings of the study the following recommendations were made to different stake holders.

1. To respective hospitals: Minimum requirements for good dispensing practice should be made available in adequate quantities for pharmacy service by taking administrative actions, Providing on job training to dispensers particularly on patient counseling; appropriate patient to dispenser ratio and adequate APTS support should be provided for transaction and service monitoring.
2. Zonal health department and regional health bureau: Continued education and training should be given to dispensers particularly on patient counseling; appropriate patient to dispenser ratio and adequate APTS support should be provided for transaction and service monitoring.
3. Other stake holders and partners: organizations working to improve patient care and rational drug use should support hospitals financially/ technically to standardize dispensing practice environment

## Acknowledgement

We wish to give Glory and praise to the Almighty God who gave us good health, strength, courage and commitment to complete the thesis. We would like to express my grateful heartfelt appreciation to hospitals in Gamo Gofa Zone, data collectors and supervisors participated in the study. Arbaminch College of Health Sciences research and publication core process. Lastly our special gratitude go to the participants of the study who shared their time and gives their genuine responses.

## Contribution of researchers

***Mende Mensa*** is senior researcher who analyzed and interpreted the findings of this study and he also prepared this document for publication. ***Akililu Ayele and Biruk Wogayehu*** conceived the study and prepared the proposal and participated in data analysis and presented the work for responsible bodies.

## Abbreviations

ABC: A class, B class and C class
ABC/VEN: A class, B class and C class medicines / Vital, essential and non-essential
AMR: Anti-Microbial resistance
APTS: Auditable Pharmaceuticals Transactions and Services
ARM: Annual Review Meeting
CTA: Consumption to stock analysis
FMOH: Federal Ministry of Health
HSDP IV: Health sector development IV (2010-2015) of Ethiopia
IFRR: Internal facility report and requisition form
MAM/SAM: Moderate Acute Malnutrition and Severe Acute Malnutrition
MDG: Millennium Development Goal
MFRF: Monthly Financial Reporting form of APTS
MOFED: Ministry of Finance and Economic Development
MSRF: Monthly Service Reporting form of APTS
RFEDB: Regional Finance and Economic Development Bureau
RHB: Regional Health Bureau
SIAPS: Systems for Improved Access to Pharmaceuticals and Services
SPS: Strengthening Pharmaceutical Systems
SPSS: Statistical Package for Social Sciences
SSA: Stock Status Analysis
STI: Sexually Transmitted diseases
STR: Stock Turnover Ratio
UN: United Nations
UNICEF: United Nations Children Fund
VEN: Vital essential and nonessential
WHO: World Health Organization

